# RBM39 alters phosphorylation of c-Jun and binds to viral RNA to promote PRRSV proliferation

**DOI:** 10.1101/2020.11.13.382531

**Authors:** Yinna Song, Yanyu Guo, Xiaoyang Li, Ruiqi Sun, Min Zhu, Jingxuan Shi, Lilin Zhang, Jinhai Huang

## Abstract

As transcriptional co-activator of AP-1/Jun, estrogen receptors and NF-κB, nuclear protein RBM39 also involves in precursor mRNA (pre-mRNA) splicing. Porcine reproductive and respiratory syndrome virus (PRRSV) causes sow reproductive disorders and piglet respiratory diseases, which resulted in serious economic losses worldwide. In this study, the up-regulated expression of RBM39 and down-regulated of inflammatory cytokines (TNF, IL-1β) were determined in PRRSV-infected 3D4/21 cells, and accompanied with the PRRSV proliferation. The roles of RBM39 altering phosphorylation of c-Jun to inhibit the AP-1 pathway to promote PRRSV proliferation were further verified. In addition, the nucleocytoplasmic translocation of RBM39 and c-Jun from nucleus to cytoplasm were enhanced in PRRSV-infected cells. The three RRM domain of RBM39 are crucial to support the proliferation of PRRSV. several PRRSV RNA (nsp4, nsp5, nsp11 and N) binding with RBM39 were determined, which may also contribute to the PRRSV proliferation. Our results revealed a complex mechanism of RBM39 by altering c-Jun phosphorylation and nucleocytoplasmic translocation, and regulating binding of RBM39 with viral RNA to prompt PRRSV proliferation. The results provide new viewpoints to understand the immune escape mechanism of PRRSV infection.

## Introduction

RNA-binding proteins (RBPs) have been found in all living organisms(1). Through binding RNAs, RBPs assemble in ribonucleoprotein complexes, which dictate the fate and the function of virtually every cellular RNA molecules. RNA-binding proteins (RBPs) can bind single-stranded or double-stranded RNAs, and play an important role in regulating RNA metabolism and gene expression as post-transcriptional regulators(2). RBPs are involved in biological processes such as RNA transcription, editing, splicing, transportation and positioning, stability and translation(3). With the extensive reports on the post-transcriptional regulatory mechanism of RNA-binding proteins, more and more scholars are focusing on their functions. The target RNA of RBPs is variable, they can bind to different region of mRNA [such as exons, introns, untranslated regions (UTRs)], or interact with other types of RNA, including non-coding RNAs(4), microRNAs, small interference RNAs (siRNA), t-RNAs, small nucleolar RNA (snoR-NA), telomerase RNA, conjugant small nuclear RNA (snRNA) and the RNA part of signal recognition particles (SRP RNA or 7SL RNA(5–7). These non-coding RNAs form a wide range of secondary structures, which combine with RBP and regulate processes such as RNA splicing, RNA modification(8), protein localization, translation, and maintenance of chromosome stability(9, 10). The functional effects of conventional RBPs depend on the target RNA-RNP complex formation. Simultaneously, the RNP complex helps RNA processing, translation, export and localization(10, 11).

RBM39 (RNA binding motif protein 39), also called CAPER, HCC1.3/1.4、Caperα 、 FSAP59 、 RNPC2(12), is a nuclear protein that is involved in precursor mRNA (pre-mRNA) splicing(13). RBM39 was first identified as an auto-antigen in a hepatocellular carcinoma patient(14). In addition, RBM39 is a transcriptional co-activator of activating protein-1 (AP-1/Jun) and estrogen receptors (eg, steroid nuclear receptors ESR1/ER-alpha and ESR2/ER-β) and NF-κB(13, 15), and its function has been associated with malignant progression in a number of cancers(14). Previous studies have reported RBM39 as a proto-oncogene with important roles in the development and progression of multiple types of malignancies(16). RBM39 is relevant to numerous precursor messenger RNA (pre-mRNA) splicing factors and RNA binding proteins. As a mRNA splicing factor, RBM39 provide regulating action of a quantity of signal pathway(17). Degradation of RBM39 can led to aberrant pre-mRNA splicing, including intron retention and exon skipping, in hundreds of gene(14). RBM39 is extremely homologous and conserved in many species (Fig. 1B). RBM39 has three RRM (RNA recognition motif) domains and the C-terminal of RRM3 domain belongs to the U2AF homology motif family (UHM), which mediate protein–protein interactions through a short tryptophan-containing peptide known as the UHM-ligand motif (ULM). However, RBM39 impacting virus proliferation and its role in innate immunity are largely unknown.

**Figure 1.**
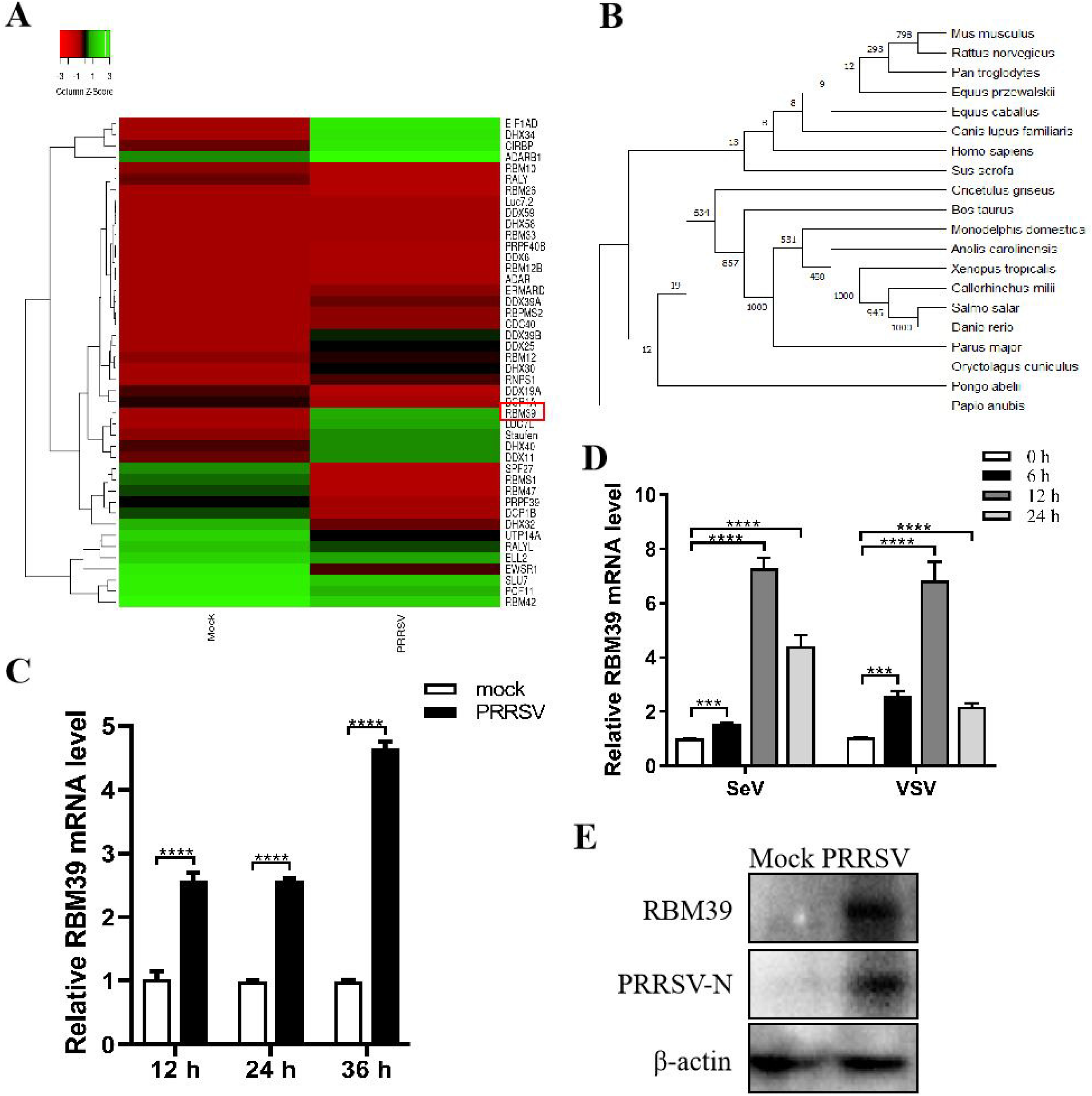
Upregulation of RBM39 in 3D4/21 cells by PRRSV infection. (A) Heat map of RNA binding protein differentially expressed genes made using online websites (http://www.heatmapper.ca/). (B) RBM39 Evolutionary tree among different species. (C) 3D4/21 cells were subjected to mock infection or infected with PRRSV at a multiplicity of infection (MOI) of 0.4. Cells were collected at the three time points (12, 24, 36 h) and subjected to real-time RT-PCR to analyze the expression level of RBM39. (D) 3D4/21 cells were infected with SeV and VSV. Then, cells were collected and extracted the total RNA of the cells at 0, 6, 12, and 24 h after infection, and analyzed the mRNA level of RBM39 by qRT-PCR. (E) 3D4/21 cell subjected to mock infection or infected with PRRSV. Cell lysates were collected at 24 h postinfection and subjected to Western blot analysis with antibody to RBM39 to analyze the protein expression. * P < 0.05; ** P < 0.01; *** P < 0.001; ****p<0.0001 (unpaired t tests). Data are representative of results from three independent experiments.

The activation protein-1 (AP-1) transcription factors are immediate early response genes involved in a diverse set of transcriptional regulatory processes(18, 19). The AP-1 complex consists of a heterodimer of a Fos family member and a Jun family member(20). The Fos family contains four proteins (c-Fos, Fos-B, Fra-1, and Fra-2), whereas the Jun family is composed of three proteins (c-Jun, Jun-B, and Jun-D)(18). The c-Jun is a central component of all AP-1 complexes and members of the basic region-leucine zipper (bZIP) family of sequence-specific dimeric DNA-binding proteins. In addition, the c-Jun is a transcription factor that recognizes and AP-1 complex binds to the enhancer heptamer motif 5’-TGA[CG]TCA-3’ (termed AP-1 sites) found in a variety of promoters(13, 18). The c-Jun gene is expressed in many cell types at low levels, and its expression is in elevated response to many stimuli, including growth factors, cytokines, foreign materials and viral infection. Some stimuli of cells result in activation of the JNK (c-Jun N-terminal kinase) and p38 groups of MAPKs(8). The c-Jun were phosphorylated at two positive regulatory sites (serine 63 and 73) residing within its amino-terminal activation domain by JNKs(18).

Porcine reproductive and respiratory syndrome virus (PRRSV) is a critical pathogen in pig and can trigger a serious negative impact on the economic development of pigs(21), which mainly causes sow reproductive disorders and piglets respiratory diseases, thus resulting in serious economic losses in the world(22). It is a variety of capsule single-stranded positive-chain virus of arteritis virus(23). Infections by PRRSV often result in increased expression of multiple genes and delayed, low-level induction of antiviral cytokines, thereby destroying the early endogenous immune response(24). The 15kb PRRSV genome is expressed through a set of subgenomic mRNA transcripts,each used for the translation of one or two open reading frames (ORFs)(25, 26). The full-length viral RNA is used for the translation of pp1a and pp1ab polyproteins(27), which are processed by viral proteases to release 14 non-structural proteins(28), including four proteases (NSP1α, NSP1β, NSP2 and NSP4), the RNA-dependent RNA polymerase (NSP9), a helicase (NSP10) and an endonuclease (NSP11)(26, 29). In this study, the RBM39 participated in PRRSV mRNA binding and viral proliferation were investigated. The dual regulation effects of RBM39 compete c-Jun to down-regulated IFN production and prompt PRRSV proliferation by stabilizing and binding with viral RNA were illustrated.

## Materials and methods

### Cell and Virus cultures

Human embryonic kidney 293 T cells (HEK293 T) and HeLa cells were cultured in Dulbecco’s modified essential medium (DMEM), Porcine alveolar macrophages (PAM) cell line 3D4/21 in RPMI-1640 medium (Biological Industries) in incubator at 37℃ with 5% CO_2_. All media were obtained from Biological Industries and supplemented with 10 % fetal bovine serum (FBS), 100 U/ml penicillin, 100μg/ ml streptomycin. The PRRSV (Porcine Reproductive and Respiratory Syndrome Virus) was preserved in our laboratory and produced by transfection of 3D4/21 cells with the PRRSV-JXwn06 infectious clone plasmid. In this study, the PRRSV was used with a titer of 10^4^ PFU/ml. Sendai virus (SeV) and vesicular stomatitis virus (VSV) was preserved in our laboratory.

### Antibodies and reagents

Murine RBM39 pAb was prepared by immunizing mice with the purified protein. Anti-PRRSV Nsp2 was a gift from China Agricultural University and Rabbit anti-PRRSV-N protein pAb was purchased from YBio Technology (YB-23941R). Mouse anti-Flag/β-actin/GADPH mAb and Peroxidase-Conjugated Goat Anti-Rabbit/Mouse IgG (H+L) were purchased from Yeasen Technology. Rabbit anti-phospho-JNK1+2+3 (Thr183 + Tyr185) pAb was purchased from Bioss (bs-1640R); Rabbit anti-c-Jun mAb was purchased from Abways Technology (CY5290); Rabbit anti-phospho-c-Jun (Ser73) mAb was purchased from Cell Signaling Technology (3270T); Rabbit anti-HA pAb, Goat anti-Mouse IgG (H+ L) Alexa Fluor 555 and Goat anti-Mouse/Rabbit IgG (H+L) FITC was purchased from Invitrogen; Mouse anti-Flag/HA-tags beads was purchased from Abmart. Phosphatase inhibitors were purchased from APE×BIO. Quick CIP were purchased from Biolabs.

### Plasmids

RBM39 and c-Jun gene was cloned from 3D4/21 cells cDNA as template. Flag/HA/Myc-RBM39 and Flag/HA/Myc-c-Jun plasmids were constructed by seamless cloning technology. Flag-RRM1, RRM2, RRM3, ΔRRM1, ΔRRM2, Δ RRM3, ΔRRM, ΔNLS were constructed by reverse amplification by Flag-RBM39 as template. Plasmid pET-28a-RBM39-X7 that delete the complete 5-terminal NLS sequence was also constructed by seamless cloning method, in order to purify the RBM39 protein and make its antibodies. AP-1/IFN-β/ Renilla luciferase reporter plasmids were constructed as previously described (23). The primers used above are shown in the table 1.

**Table 1.**
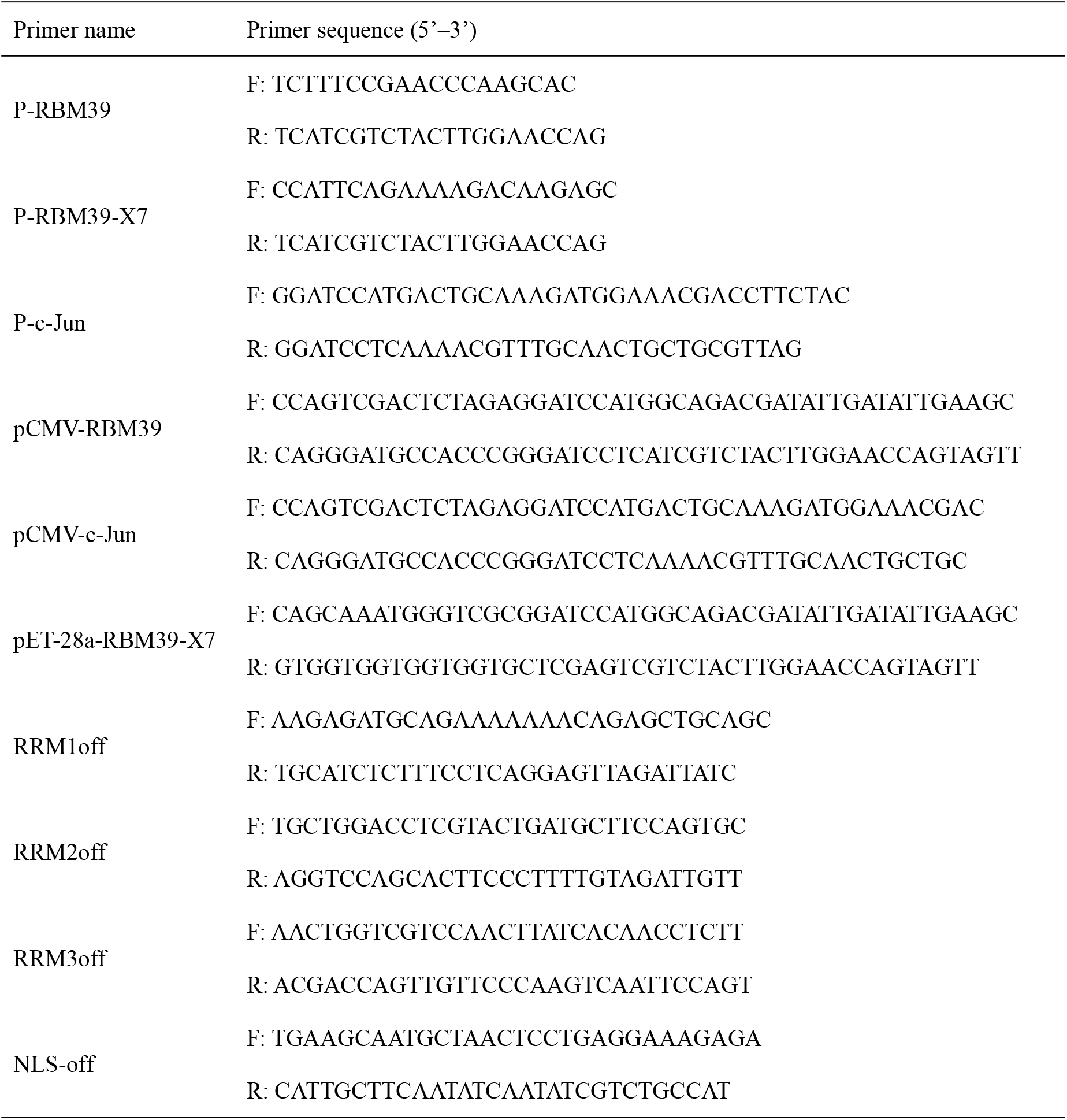
Primers used in PCR amplification.

### Transient transfection of eukaryotic expression plasmid into cells

Previous to transient transfection, adherent HEK293T, HeLa or 3D4/21 cells were seeded in a 6/12/24-well plate. Cells were transfected at appropriate confluence with eukaryotic expression plasmid. Briefly, 10 μM polyethyleneimine was preheated at 85 degrees and replace the old medium with Reduced-Serum Medium (Opti-MEM, Thermo Fisher Scientific) before transfection 1 h in advance. Select an appropriate mixing ratio (3:1/PEI: DNA quality) to transfect cells. Add a suitable equal volume of medium to the two centrifuge tubes, then add the appropriate plasmid and PEI, shake and mix, and place at room temperature for 5 minutes. Mix the two together and place at room temperature for 20-30 minutes. Afterwards, add the mixture to the cells and incubate the cells in a 37°C incubator containing 5% CO2. After 4-6 hours, discard the mixed solution and replace to complete medium, and continue to grow cells for 24-48 h.

### Western blot

Cell samples were lysed by radioimmunoprecipitation assay (RIPA) Lysis Buffer (Solarbio) added with protease inhibitor cocktail phenylmethanesulfonyl fluoride (PMSF, Solarbio). Cells lysis supernatants were added to 5x Loading Buffer, following boiled for 10 min and separated by SDS-PAGE. The separated proteins were transferred onto a PVDF blotting membrane (GE Healthcare) and blocked for 1h with 5% skimmed milk in 1×TBST (Tris-buffered saline containing 0.05% Tween-20) at room temperature, followed by incubation with the primary antibodies diluted in 1×TBST at 4°C overnight and secondary HRP-conjugated antibodies diluted in 1×TBST for 1h at room temperature. Finally, the membrane was washed with 1×TBST three times and detected through Pierce ECL Western Blotting Substrate (Thermo Scientific, Waltham, MA,USA) and exposed through Chemi Doc XRS Imaging System (BIO-RAD, USA).

### Quantitative real-time PCR

Quantification of the relative levels of gene expression was performed using qRT-PCR. Total RNAs were extracted by RNAiso plus (Takara). The cDNA was synthesized with EasyScript First-Strand cDNA Synthesis SuperMix according to the manufacturer’s instructions (TransGene). The relative levels of gene expression were analyzed by ABI 7500 real-time PCR system using TransStart Top Green qPCR SuperMix (TransGene) with three-step amplification and the data were calculated by the comparative cycle threshold (C_T_) method (2^−ΔΔCt^). The thermal cycler program consisted of 95°C for 10 min and then 45 cycles at 95°C for 15 s, 58°C for 30 s, and 72°C for 30 s. The β-actin was used as an internal control for normalization. All the primers used for quantitative real-time PCR are shown in Table 2.

**Table 2.**
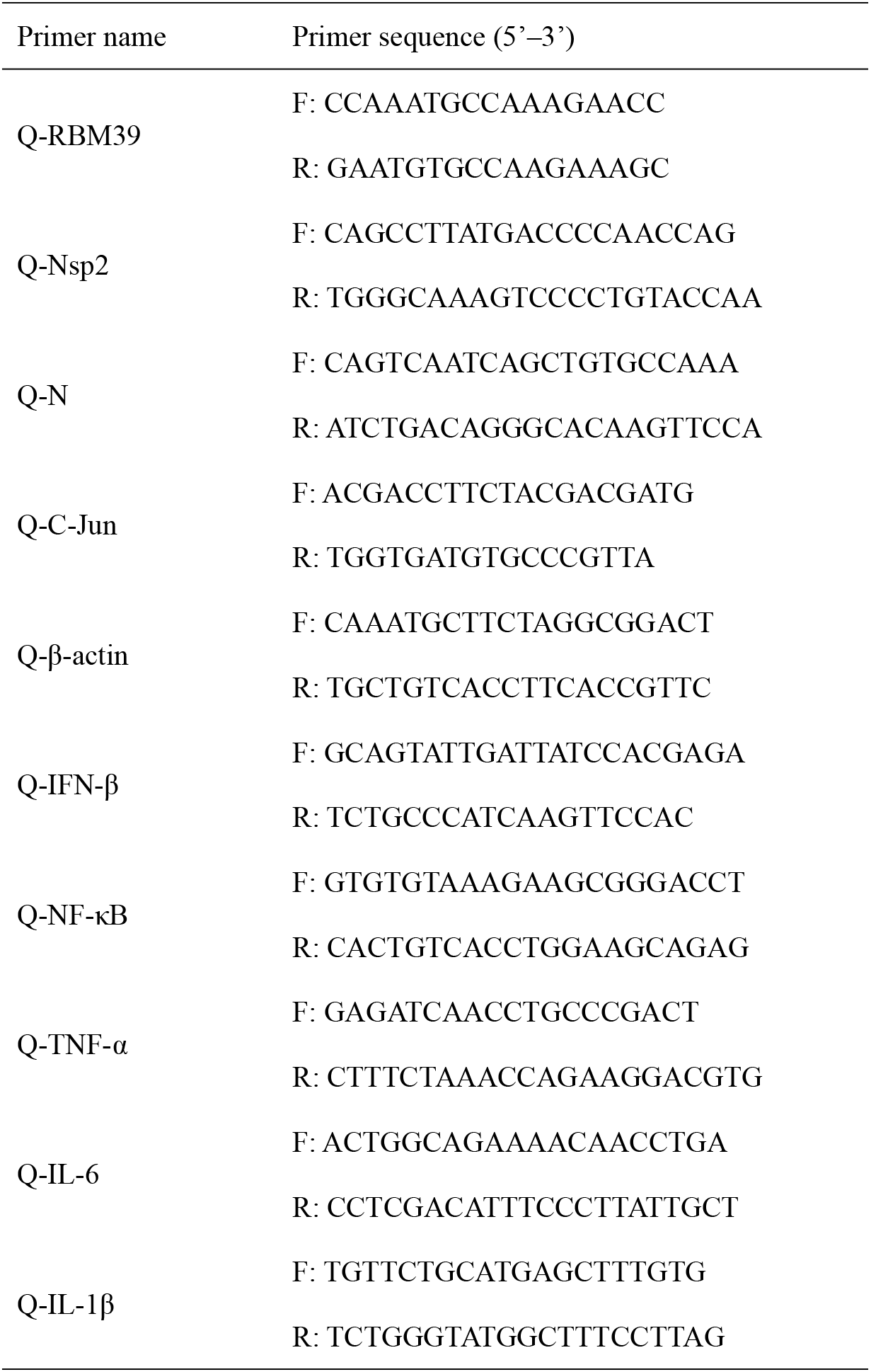
Primers used in quantitative real-time PCR.

### Dual luciferase reporter assay

HEK293 T cells plated to 24-well plates at 70%-80% confluence were transfected with RBM39, AP-1 luciferase reporter plasmid, PRRSV-JXwn06 infectious clone plasmid, siRBM39 or sic-Jun (Sigma) and the Renilla luciferase control reporter plasmid was used as a reference. The cells were collected and lysed at 24 h post-transfection, and the cell supernatant was used to detect luciferase activity through Dual Luciferase Reporter Gene Assay Kit (Yeasen Technology) according to the manufacturer’s instructions.

### Immunoprecipitation (IP) and co-immunoprecipitation (co-IP) assays

HEK293 T Cells cultivated in six-well plates were transfected or co-transfected with appropriate eukaryotic expression plasmid. At 24 h posttransfection, the cells were harvested and resuspended by PBS, and lysed on ice for 15 min in 400 μl RIPA lysis buffer. Then the samples were centrifuged at 12,000 rpm for 5 min. We took out the 50 μL supernatant in new tube, added 10 μL loading buffer and boiled them to make input samples. Subsequently, anti-Flag-labeled beads (Sigma) were activated by RIPA and added the rest of cell lysis supernatant. The reaction mixture was incubated at 4°C overnight. The beads were washed three times with lysis buffer containing PMSF for 5 min each time and added 50 μL lysates and 10 μL loading buffer, boiled for 10 min to make IP samples. Finally, the proteins bound to the beads were separated via SDS-PAGE, transferred to polyvinylidene fluoride (PVDF) membrane and detected with the proper antibodies.

### Indirect immunofluorescent assay (IFA) and confocal microscopy

Adherent HEK293T, HeLa or 3D4/21 cells grown on glass coverslips in 12-well plates (Corning Inc., Corning, NY, USA) at 30% to 50% confluence were transfected with plasmids expressing RBM39 and/or c-Jun. The empty vector plasmid was used as a negative control (NC). Subsequently, 3D4/21 cells were infected with PRRSV at an MOI of 0.4. At 24 h posttransfection or infection, the cells washed with 1 × phosphate-buffered saline (PBS), fixed with 4% paraformaldehyde for 30 min at room temperature, permeabilized with 0.5% Triton X-100 for 15 min and blocked with 5% bovine serum albumin (BSA) dissolved in 1 × PBS containing 0.05% of Tween-20 (PBST) for 2h at room temperature. Anti-HA (hemagglutinin), anti-Flag and anti-PRRSV-N primary antibodies were diluted in 2% BSA and incubated with slides overnight at 4°C. Following a wash performed with PBS 3 times for 5 min each time, the cells were incubated with fluorescein isothiocyanate (FITC)-conjugated anti-mouse/rabbit IgG or IF555-conjugated anti-mouse IgG secondary antibody diluted in 2% BSA at 37°C for 1 h in a humidified chamber. The slides were washed with PBST and nuclei were counterstained with Hoechst 33258 dye (Solarbio) for 5 min. Confocal images were collected with confocal laser scanning microscope (UltraView Vox, PerkingElmer) and taken at 40 × or 100 × magnification (Olympus).

### RNA interference

A small interfering RNA (siRNA) assay was performed through siRNA targeting the knockdown of RBM39 or c-Jun gene (siRBM39 or sic-Jun) and a negative control (NC), which were synthesized from Sigma (Table 3). For siRNA transfection, 3D4/21 cells plated in 12-well plates and grown to 70%-80% confluence were transfected with 50 nmol of siRNAs using 10 μM polyethyleneimine. 24 h after transfection, cells were infected with PRRSV at an MOI of 0.4, and the cells were incubated for additional 24 h. Subsequently, cells were collected for qRT-PCR and Western blotting analyses to confirm the expression levels.

**Table 3.**
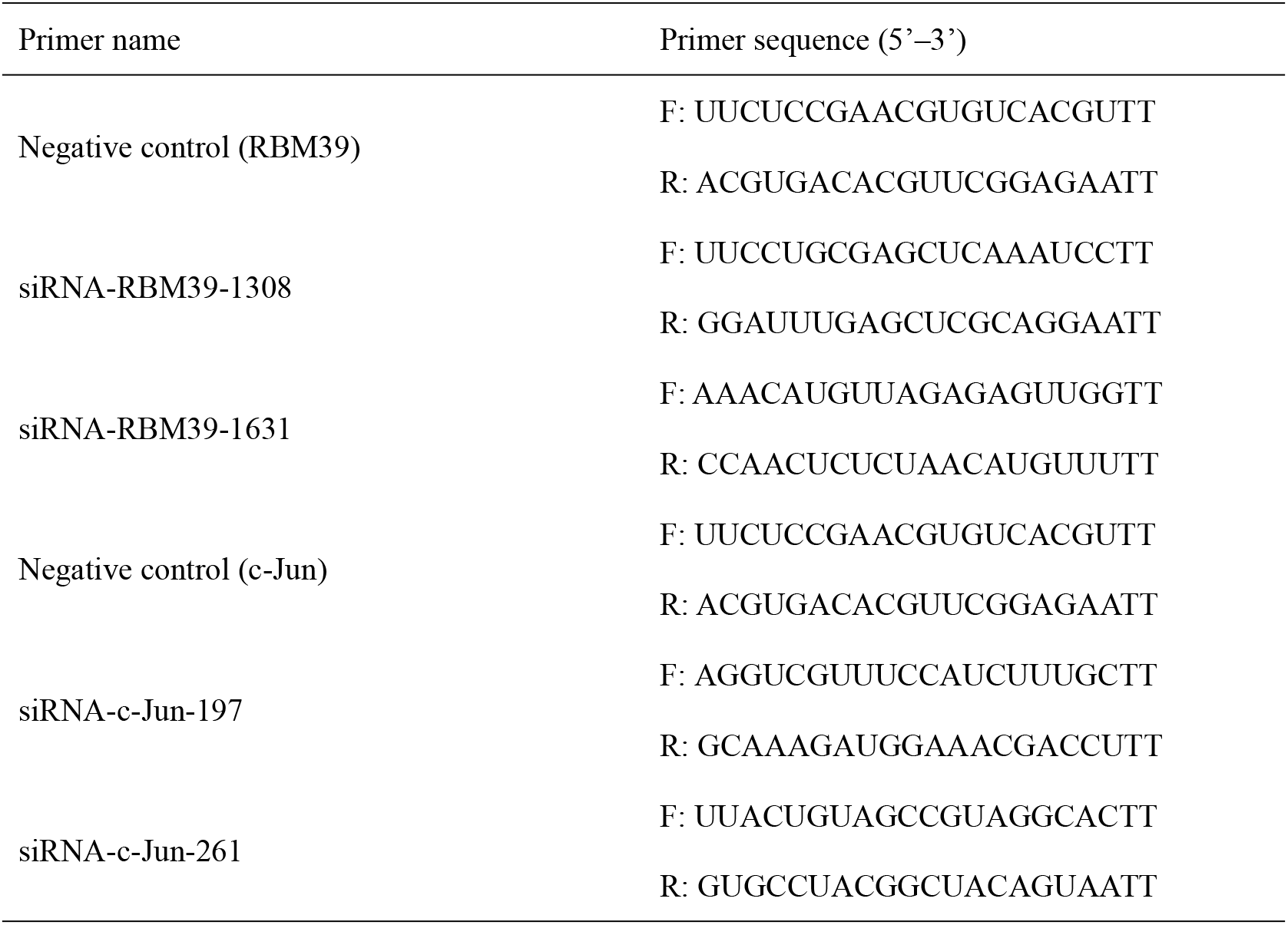
Primers used in RNA interference assay.

### RNA immunoprecipitation (RIP)

3D4/21 cells plated in 6-well plates and grown to 70%-80% confluence were transfected with 2μg of HA-RBM39 or Flag-RBM39 plasmids. Then the cells were infected with PRRSV 12 h after transfection. After another 24 hours, the cells were collected and lysed with 400 μl RIPA supplemented with murine RNase inhibitor (1:100). Part of the supernatant was used for immunoblotting analysis, and the remaining supernatant was incubated with HA/Flag-tags beads overnight at 4℃. The co-precipitated RNAs were extracted and detected through reverse transcription PCR via corresponding primers.

### Statistical analysis

Data were presented as the mean of three independent replicates. Statistical significance was analyzed by the paired two-tailed t-test using Origin and GraphPad Prism software. Statistical differences were considered to be statistically significant at P-value < 0.05 (*p < 0.05; **p < 0.01; ***p < 0.001; ****p<0.0001).

## Results

### Upregulation of RBM39 in 3D4/21 cells by PRRSV infection

PRRSV infection can result to upregulation or downregulation of many genes in cells.In order to find out genes that displayed significant changes in transcription after PRRSV infection, a transcriptome sequencing analysis derived from 3D4/21 cell was performed. RBM39 was found to be significantly upregulated after PRRSV infection by statistical analysis (Fig. 1A). To investigate the levels of RBM39 accumulation during viral infection, we infected 3D4/21 cells with PRRSV at a multiplicity of infection (MOI) of 0.4. After Collecting cells and extracting total RNA at different time points (12 h, 24 h and 36 h) postinfection, mRNA abundance of RBM39 was analyzed by quantitative reverse transcription-PCR (qRT-PCR). Compared with the control group, the mRNA levels of RBM39 significantly increased at three different time points after PRRSV infection (Fig. 1C). In order to verify the expression level of RBM39 after infection with other RNA viruses, we infected 3D4/21 cells with SeV and VSV, and then extracted the total RNA of the cells at 0, 6, 12, and 24 h after infection, and analyzed the mRNA level of RBM39 by qRT-PCR. The results showed that RBM39 mRNA levels increased significantly at four time points (Fig. 1D). Furthermore, to analyze the protein expression at the translation level, cell lysates were collected at 24 h postinfection and subjected to Western blot analysis with antibody to RBM39. The results showed that the protein expression in cellular level of RBM39 after PRRSV infection was dramatic increase compared with control groups (Fig. 1E).

### RBM39 contributes to PRRSV proliferation

The effect of RBM39 on the virus has not been reported before. In order to explore the effect of RBM39 on PRRSV proliferation, Flag-RBM39 expression plasmids were transfected into 3D4/21 cells. Total RNA was extracted from the cells at 24 h posttransfection, and subsequently, transcription levels of PRRSV NSP2 and N gene were analyzed by qRT-PCR. Compared with the control groups, the overexpression of RBM39 promoted the mRNA level of the PRRSV NSP2 and N gene (Fig. 2A). To figure out the effect of RBM39 on viral proliferation at protein levels, we collected cells at five different time point (6, 12, 24, 36, 48 hs) after PRRSV infection respectively, and subjected to Western blot analysis with antibody to NSP2. The results of Western blot analysis revealed that the expression level of NSP2 increased with time after transfection of RBM39 (Fig. 2D).

**Figure 2.**
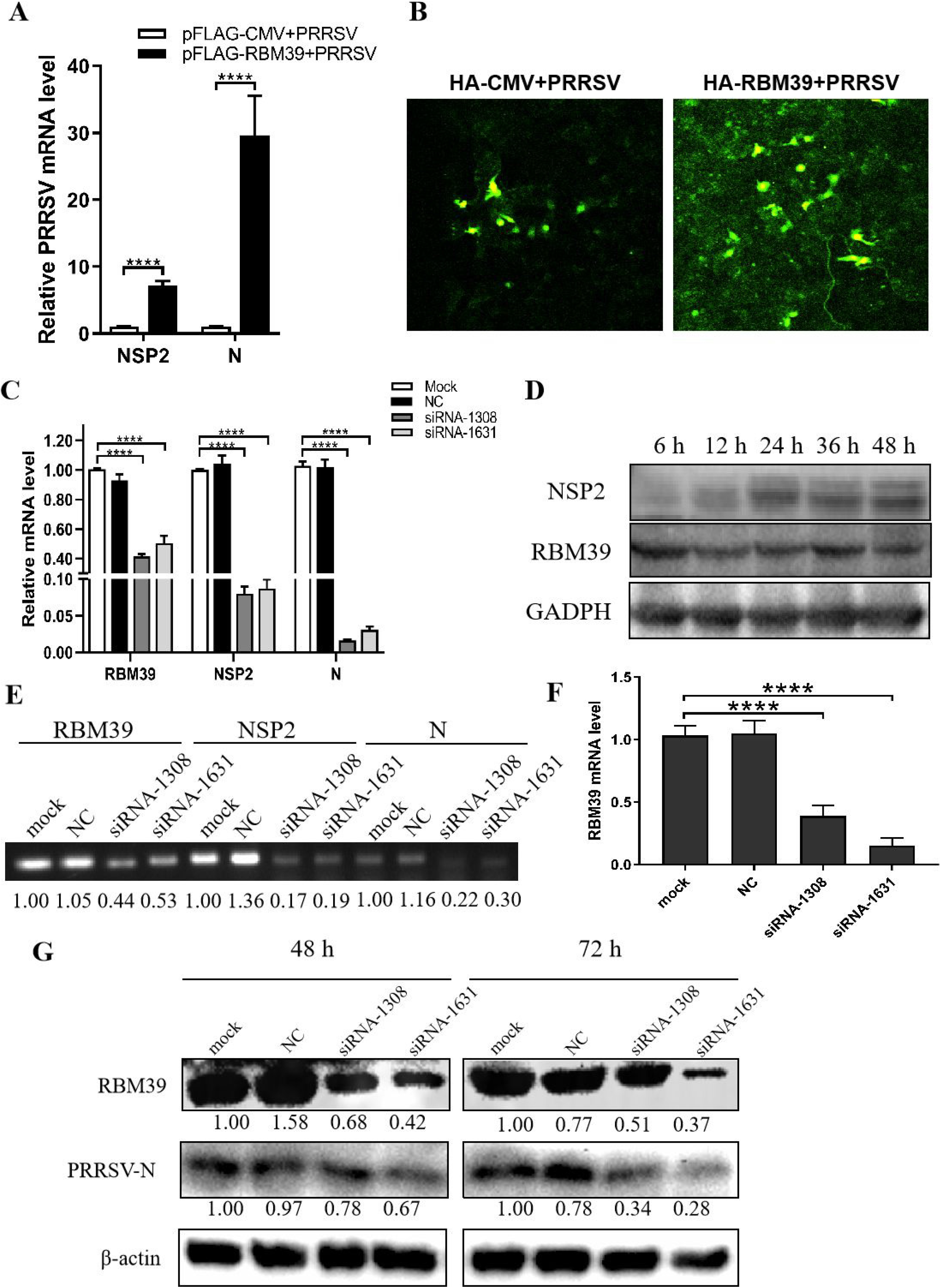
RBM39 contributes to PRRSV proliferation. (A, B and D) 3D4/21 cells were transfected with Flag/HA-RBM39 or empty vector and infected with PRRSV at 12 h posttransfection. The infected cells were collected at different time piont (24 h (A) or 6, 12, 24, 36, 48 h (D)) postinfection. The mRNA loads of PRRSV N and Nsp2 were tested by qRT-PCR (A), and protein expression of PRRSV N were tested by WB (D). (B) The cell slide was incubated overnight at 4° C with the primary antibody of PRRSV N and then was incubated for one hour at room temperature with FITC fluorescent secondary antibody. The expression of PRRSV N was detected by Laser confocal fluorescence microscope. (C, E, F and G) 3D4/21 cells were transfected with siRNA of RBM39 or negative control (NC) or were left untransfected. 12 h after transfection, cells were infected with PRRSV. (C and F) The cells were collected at 24 h postinfection. The mRNA loads of RBM39, PRRSV N and Nsp2 were tested by qRT-PCR. (E) The agarose gel electrophoresis (AGE) of RBM39 PRRSV N and Nsp2 after qRT-PCR. (G) RBM39 and PRRSV protein expression levels were examined by WB after 48 or 72 h. * P < 0.05; ** P < 0.01; *** P < 0.001; ****p<0.0001 (unpaired t tests). Data are representative of results from three independent experiments.

Subsequently, we designed RBM39 small interfering RNA (siRNA) that targeted RBM39 gene and inhibited the transcription and protein expression of RBM39 in order to further clarify the effect of RBM39 on PRRSV proliferation. Consistent with previous results, knockdown of RBM39 reduced the PRRSV proliferation to some extent at the transcription and protein levels (Fig. 2C, E and G). To further verify the enhancement of RBM39 on PRRSV, we used immunofluorescence to detect the expression of PRRSV N after Flag-RBM39 transfected into 3D4/21 cells. Compared with the empty vector group, the RBM39 overexpression group can observe more green fluorescent signals in the field of view (Fig. 2B). The results indicate that changes in the transcription and translation levels of RBM39 can affect PRRSV proliferation and that RBM39 contributes to PRRSV proliferation.

### The RRM domain of RBM39 is crucial to PRRSV proliferation

In order to determine the effect of the three RRM domains of RBM39 on the proliferation of PRRSV, we constructed 8 truncated forms of RBM39 via the reverse amplification method, namely Flag-RRM1, RRM2, RRM3, ΔRRM1, ΔRRM2, ΔRRM3, ΔRRM, ΔNLS (Fig. 3A). It is worth noting that the DNA sequence that connects each domain is preserved. 3D4/21 cells were transfected with these 8 truncations and infected with the virus. After 24 hours, the proliferation of PRRSV was detected by qPCR and Western Blot analysis. The results of qPCR and Western Blot analysis showed that, compared with the control group, the proliferation of PRRSV in the experimental group transfected with Flag-RRM1, RRM2, RRM3, ΔRRM1, ΔRRM2, ΔRRM3, ΔRRM plasmid was inhibited (Fig. 3B-J). However, the proliferation of PRRSV in the group transfected with Flag-ΔNLS had no significant difference (Fig. 3B, J). It shows that the three RRM domains of RBM39 play an important role in promoting the proliferation of PRRSV.

**Figure 3.**
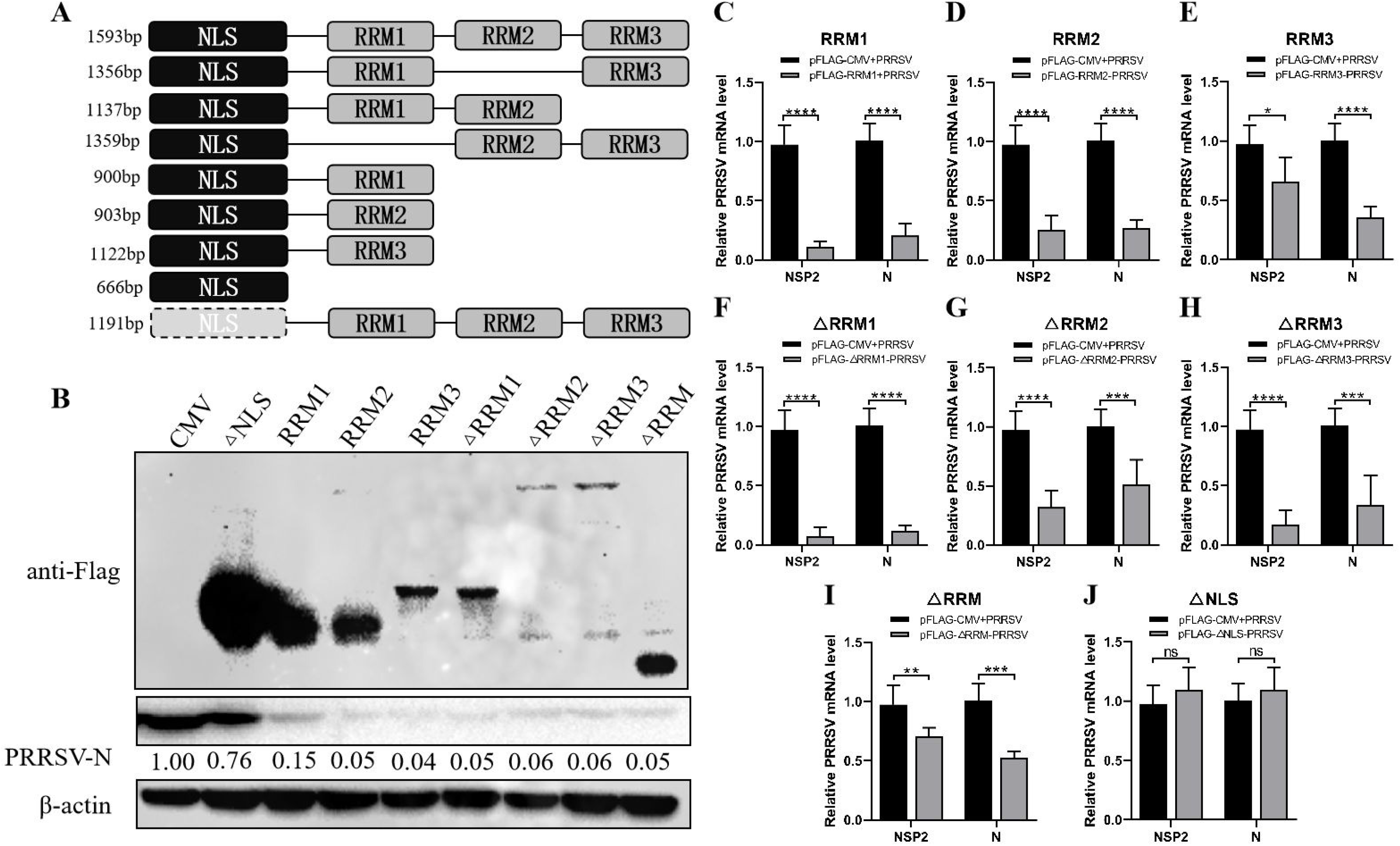
The RRM domain of RBM39 is important to promote the proliferation of PRRSV. (A) The constructed 8 truncated forms of RBM39 via the reverse amplification method, namely Flag-RRM1, RRM2, RRM3, ΔRRM1, ΔRRM2, ΔRRM3, ΔRRM, ΔNLS. (B-J) 3D4/21 cells were transfected with 8 truncations and infected with PRRSV after 12 h. After 24 hours, the proliferation of PRRSV was detected by Western Blot analysis (B) and qRT-PCR (C-J). * P < 0.05; ** P < 0.01; *** P < 0.001; ****p<0.0001 (unpaired t tests). Data are representative of results from three independent experiments.

### The promotion of PRRSV proliferation by RBM39 is related to innate immunity

Previous studies showed that RBM39 is beneficial to PRRSV proliferation. To test the negative regulatory of RBM39 effect on innate immunity, we examined the mRNA levels of some transcription factors and cytokines related to innate immunity, such as NF-κB, TNF-α, IFN-β, IL-1β, IL-6. The results shows that overexpression of RBM39 drastically decreased the transcription levels of these gene in 3D4/21 cells (Fig. 4A). However, knockdown of RBM39 by siRNA contributed to the mRNA levels of them (Fig. 4B, D). To investigate whether RBM39 impacts IFN-β-related signaling pathway, HEK293T cells were transfected with HA-RBM39 and PRRSV infectious clone plasmid or co-transfected with them. Subsequently, we measured the activity of IFN-β promoter reporter gene and found that the activation was increased by PRRSV stimuli but reduced by RBM39 overexpression compared with the control group (Fig. 4C). Most important of all, when RBM39 and PRRSV co-transfected, the activity is lower than that of the group transfected with PRRSV alone, indicating that the overexpression of RBM39 may promote PRRSV proliferation by inhibiting the IFN signaling pathway (Fig. 4C). In conclusion, these data demonstrated that RBM39 might negatively regulate antiviral immune responses and resulting in the promotion of PRRSV proliferation.

**Figure 4.**
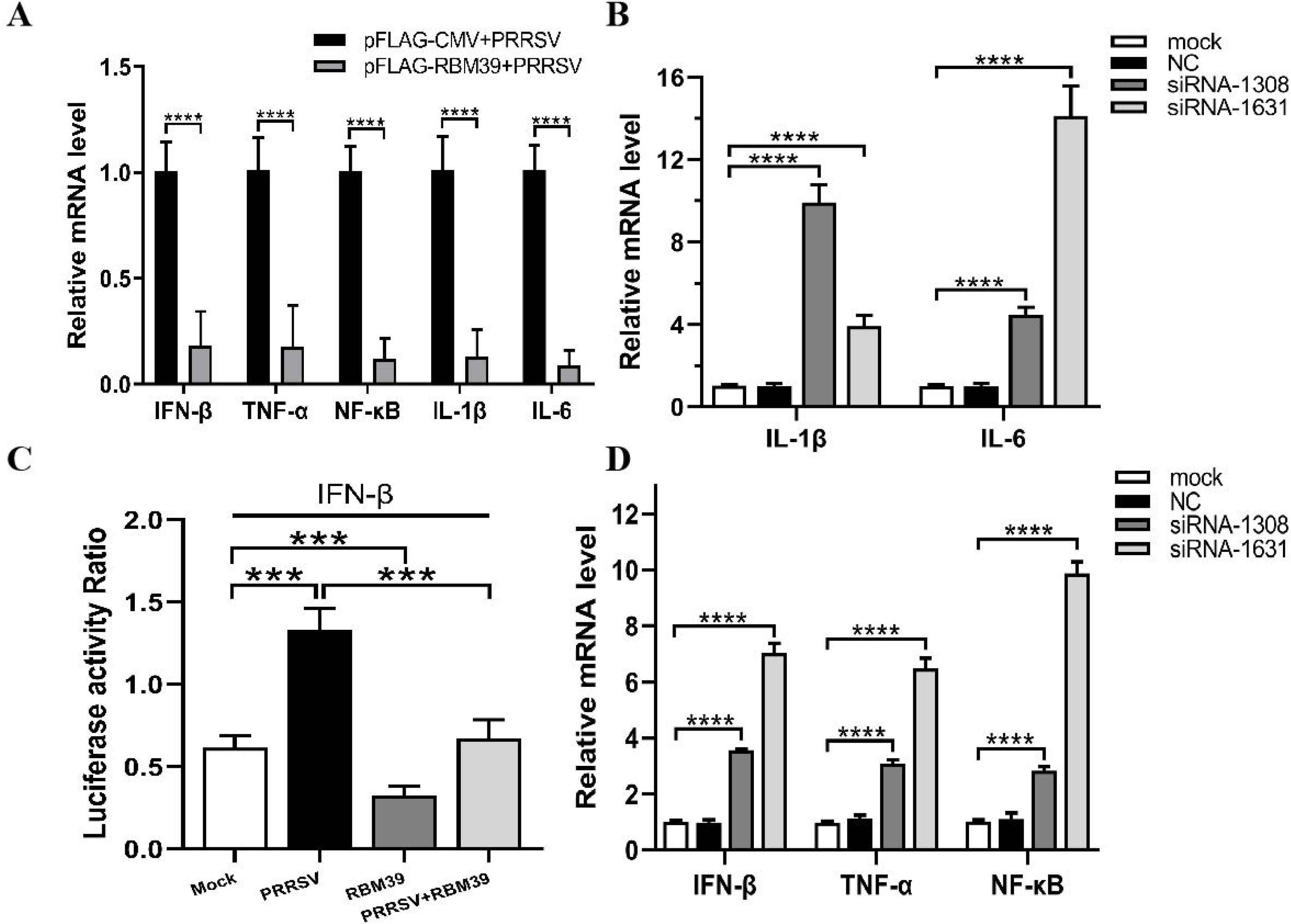
The promotion of PRRSV proliferation by RBM39 is related to innate immunity. (A) 3D4/21 cells were transfected with Flag-RBM39 or empty vector; 12 h after transfection, cells were infected with PRRSV and the mRNA levels of NF-κB, TNF-α, IFN-β, IL-1β, IL-6 were examined by qRT-PCR analysis at 12 h postinfection. (B and D) 3D4/21 cells were transfected with siRNA of RBM39 or the NC and infected with PRRSV after 12 h transfection. The mRNA levels of NF-κB, TNF-α, IFN-β, IL-1β, IL-6 were examined by qRT-PCR analysis. (C) HEK293 T cells were transfected or co-transfected with HA-RBM39 and PRRSV infectious clone plasmid. The luciferase activity of IFN-β promoter reporter gene was measured through Dual Luciferase Reporter Gene Assay Kit (Yeasen Technology). * P < 0.05; ** P < 0.01; *** P < 0.001; ****p<0.0001 (unpaired t tests). Data are representative of results from three independent experiments.

### Interaction and co-localization between RBM39 and c-Jun

Previous research and mass spectrometry (MS) results found that RBM39 binds and interact with the AP-1 component c-Jun (Fig. 5A). In addition, the data displays that the cryptic autonomous transactivation domain (AD) of RBM39 combined and interacted with the basic region-leucine zipper (bZIP) of c-Jun. In previous studies, the interaction region of RBM39 included the end of the RRM2 (RNA recognition motif 2) domain, the whole RRM3 domain, and the sequence connecting the two in the middle. Therefore, We are curious about whether RBM39 was capable of interacting with c-Jun when there is only RRM2 or RRM3. Afterwards, the plasmids pFlag-RBM39 (full-length), RRM2, RRM3, ΔRRM1, ΔRRM or empty vector and pHA-c-Jun was co-transfected into HEK293T respectively. Subsequently, cells were harvested after 24 h and interaction between RBM39 and c-Jun was detected via co-immunoprecipitation. The results shows that c-Jun can interact with full-length RBM39, RRM2, RRM3 and ΔRRM1 but not ΔRRM or empty vector (Fig. 5B). It is worth noting when both RRM2 and RRM3 domains exist, the interaction between RBM39 and c-Jun is stronger than only one of them exists (Fig. 5B). Subsequently,we co-transfected plasmids pFlag-RBM39+pHA-c-Jun or pFlag-c-Jun+pHA-RBM39 into HEK293T cell lines, and detected the co-localization of RBM39 and c-Jun. Confocal microscopy showed that both RBM39 and c-Jun expressed and co-localized in the nucleus, while similar results were not observed in the control cells (Fig. 5C-F).

**Figure 5.**
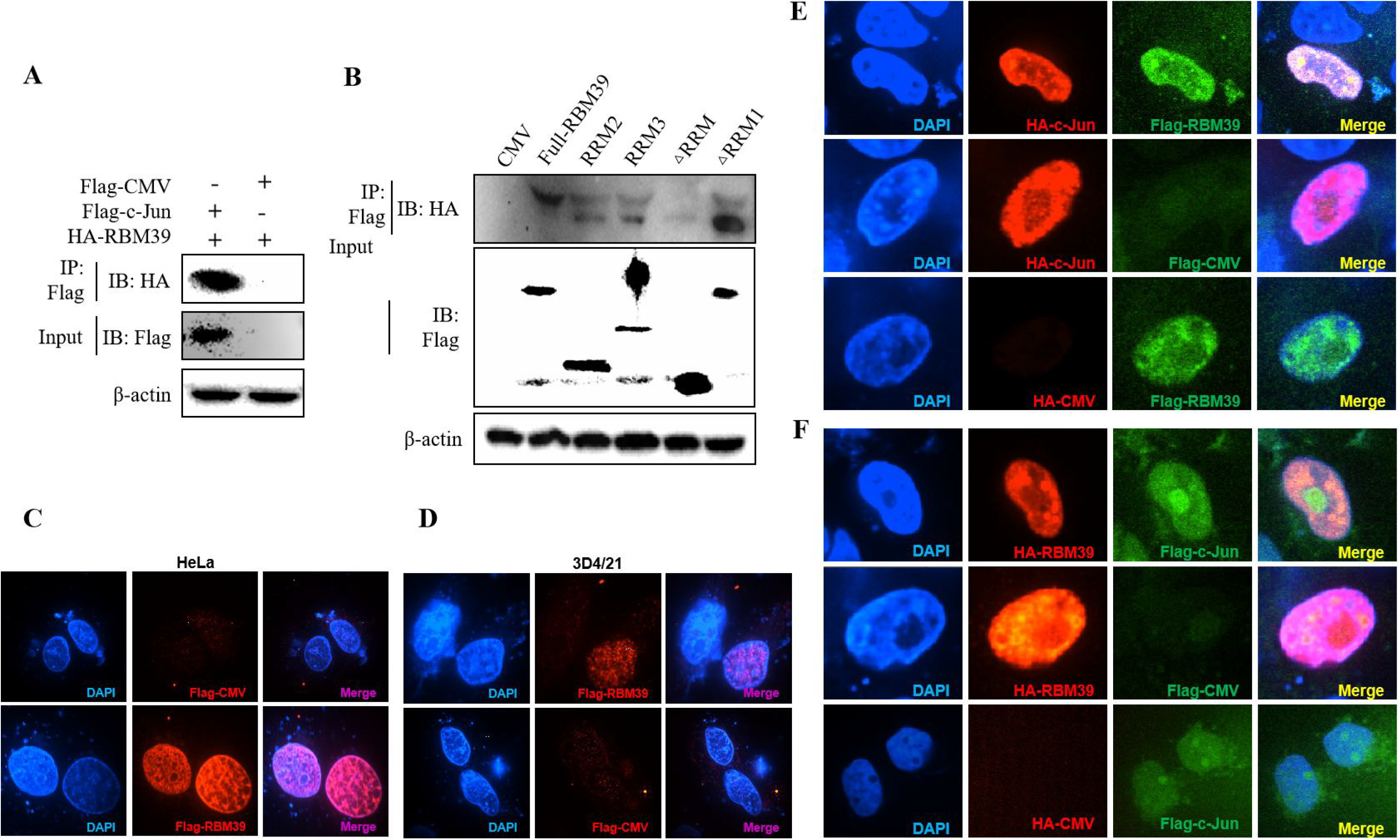
Interaction and co-localization between RBM39 and c-Jun. (A) HEK 293T cells were co-transfected with Flag-c-Jun (or empty vector) and HA-RBM39. (B) The plasmids Flag-RBM39 (full-length), RRM2, RRM3, ΔRRM1, ΔRRM or empty vector and pHA-c-Jun was co-transfected into HEK293T respectively. At 24 h after transfection, the cells lysates of (A) and (B) were precipitated with anti-Flag-labeled beads (Sigma) and were further detected by WB with anti-Flag/HA antibody. (C-F) HEK293T cell were co-transfected with pFlag-RBM39+pHA-c-Jun or pFlag-c-Jun+pHA-RBM39. After 24 h, cells were fixed and doubly stained with rabbit anti-HA mAb and mouse anti-Flag antibody followed by FITC-conjugated anti-rabbit IgG (green) or IF555-conjugated anti-mouse IgG (red). Nuclei were stained with Hoechst 33258 dye (blue). Interaction and nuclear localization of RBM39 and c-Jun were observed using Laser confocal fluorescence microscope. Scale bar: 14μm. Data are representative of results from three independent experiments.

### RBM39 down-regulates AP-1 signaling pathway

The c-Jun is an important part of AP-1 signaling pathway and a significant immune-related protein. To investigate whether RBM39 impacts AP-1 signaling pathway, HEK293 T cells were transfected or co-transfected with HA-RBM39 and AP-1 promoter reporter plasmid. The results display that the activity of AP-1 promoter reporter gene was reduced by RBM39 overexpression in a dose-dependent manner (Fig.6A, B).

**Figure 6.**
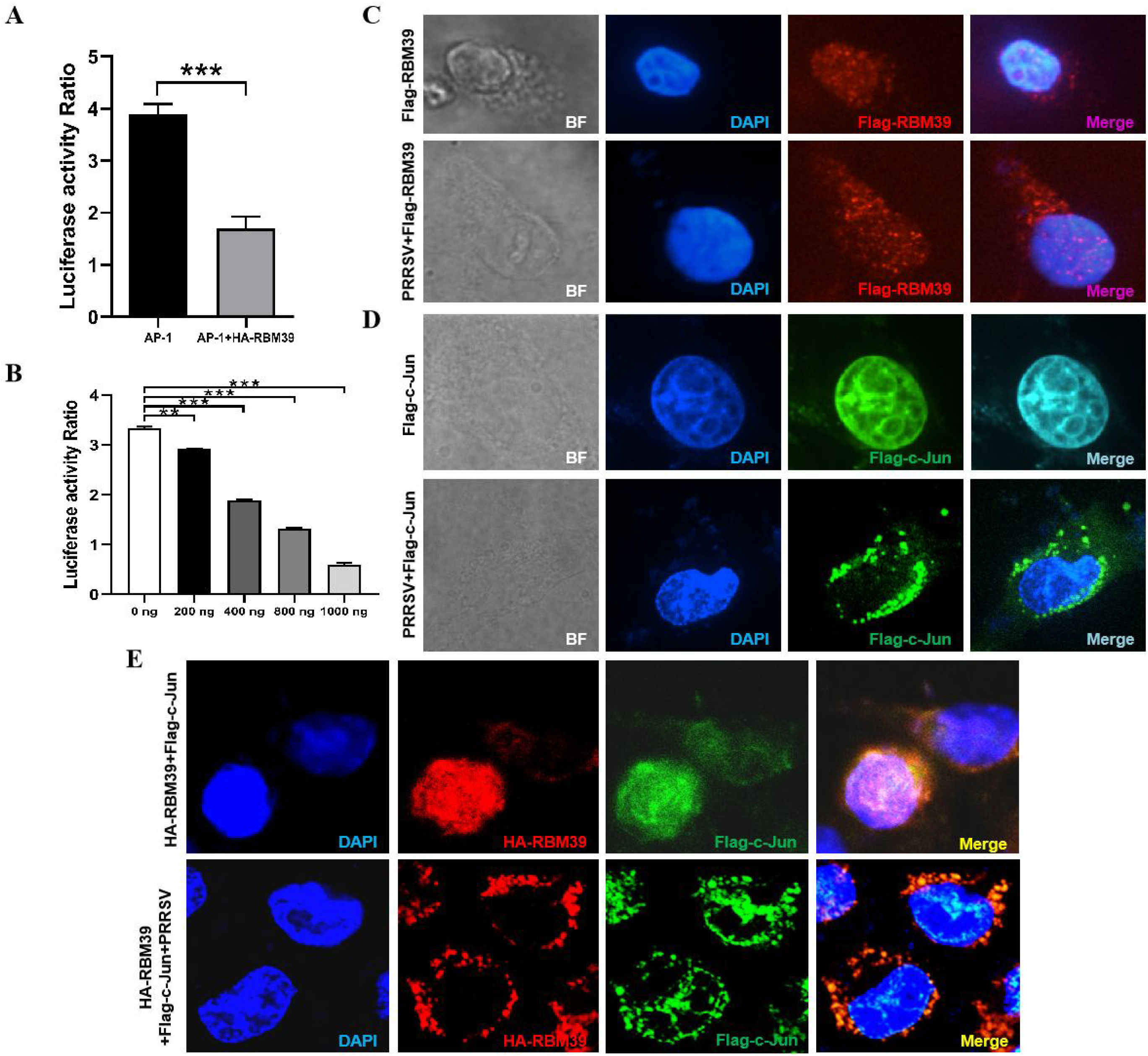
RBM39 down-regulates AP-1 signaling pathway and PRRSV infection triggers out of the nucleus of translocation of RBM39 and c-Jun. (A and B) HEK293 T cells were transfected or co-transfected with AP-1 promoter reporter plasmid and different concentrations of HA-RBM39. The luciferase activity of AP-1 promoter reporter gene was measured through Dual Luciferase Reporter Gene Assay Kit (Yeasen Technology). (C, D and F)The 3D4/21cells were respectively transfected or co-transfected with HA-RBM39 and Flag-c-Jun plasmids, 12 h after transfection, cells were infected with PRRSV at an MOI of 0.4. After 24 h, cells were fixed and doubly stained with rabbit anti-HA mAb and mouse anti-Flag antibody followed by FITC-conjugated anti-rabbit IgG (green) or IF555-conjugated anti-mouse IgG (red). Nuclei were stained with Hoechst 33258 dye (blue). Interaction and nuclear localization of RBM39 and c-Jun were observed using Laser confocal fluorescence microscope. Scale bar: 14μm. * P < 0.05; ** P < 0.01; *** P < 0.001; ****p<0.0001 (unpaired t tests). Data are representative of results from three independent experiments.

### RBM39 impair phosphorylation of c-Jun to down-regulation of AP-1 signaling pathway

To test whether RBM39 down-regulates the AP-1 pathway by affecting c-Jun, we designed siRNA to knock down the expression of c-Jun (Fig. 7A, E). First, PRRSV infection triggers the activation of the AP-1 signaling pathway (Fig. 7B, C, I), and both RBM39 overexpression and c-Jun konckdown led to a decrease in the activity of the AP-1 promoter reporter gene (Fig. 7C, F). On this basis, we hypothesized that the down-regulation of AP-1 activity by RBM39 is not due to its interaction with c-Jun, therefore, RBM39 and siRNA-c-Jun should reduce AP-1 activity to a greater degree than either. However, the results showed that RBM39 and siRNA-c-Jun did not have a synergistic effect with each other (Fig. 7F), which indirectly indicates that the down-regulation of the AP-1 pathway by RBM39 is achieved by affecting c-Jun. Similarly, the activity of AP-1 promoter reporter gene was increased by PRRSV stimuli but reduced by RBM39 and PRRSV co-transfection compared with the control group (Fig. 7B, C, I). However, the activity of AP-1 was not significantly reduced after c-Jun was knocked down by siRNA compared with the co-transfection of RBM39 and PRRSV (Fig. 7B, I), indicating that knockdown of c-Jun have not a synergistic effect on the reduction of AP-1 activity caused by RBM39 (Fig. 7I), further indicating the down-regulation of the AP-1 pathway by RBM39 is related to c-Jun.

**Figure 7.**
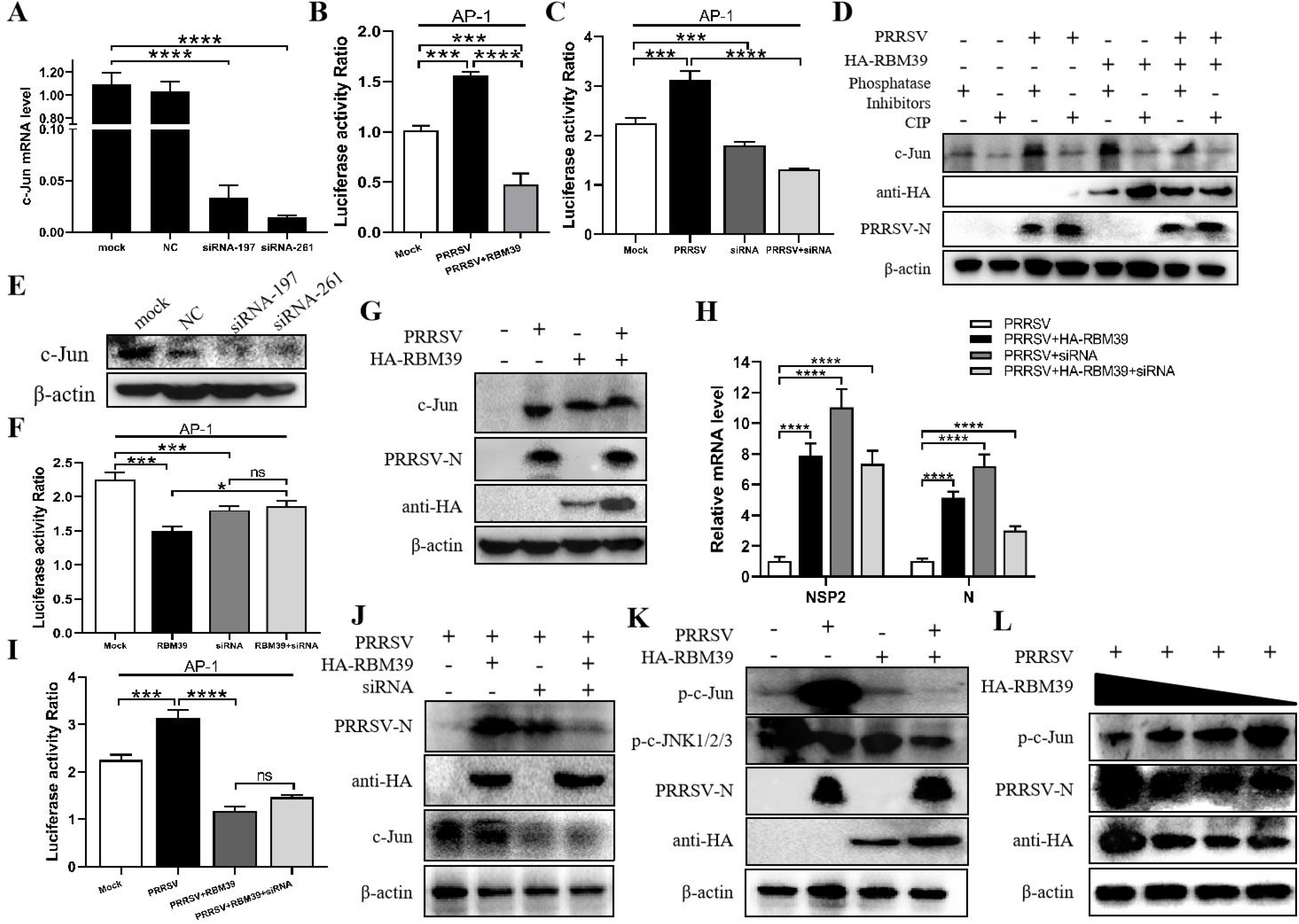
RBM39 affect phosphorylation of c-Jun to down-regulation of AP-1 signaling pathway. (A and E) 3D4/21 cells were transfected with siRNA of c-Jun or the NC, and the mRNA and protein levels of c-Jun were examined by qRT-PCR (A) and WB analysis (E). (B, C, F and I) HEK 293T cells were transfected or co-transfected with RBM39, siRNA of c-Jun, PRRSV infectious clone plasmid, AP-1/ Renilla luciferase reporter plasmids. At 24 h after transfection, the luciferase activity of AP-1 promoter reporter gene was measured through Dual Luciferase Reporter Gene Assay Kit (Yeasen Technology). (D) 3D4/21 cells were transfected or untransfected with HA-RBM39. 12 h after transfection, cells were infected or no infected with PRRSV, and phosphatase Inhibitors/CIP was used to treat cell lysate supernatant. Then cells were collected and analyzed by WB. (G) 3D4/21 cells were transfected or untransfected with RBM39. 12 h after transfection, cells were infected or no infected with PRRSV. The cells were collected and analyzed by WB after 24 h. (H and J) 3D4/21 cells were transfected or co-transfected with RBM39 and siRNA of c-Jun, respectively. 12 h after transfection, cells were infected with PRRSV. At 24 h after transfected, mRNA levels of PRRSV N and Nsp2 were tested by qRT-PCR (H), and the protein levels of PRRSV N were detected by WB (J). (K) 3D4/21 cells were transfected or untransfected with RBM39. 12 h after transfection, cells were infected or no infected with PRRSV. The cells were collected and analyzed by WB after 24 h. (K) 3D4/21 cells were transfected or untransfected with different concentrations of RBM39 (2μg, 1μg, 1.5μg, 0.5μg). 12 h after transfection, cells were infected or no infected with PRRSV. The cells were collected and analyzed by WB after 24 h. * P < 0.05; ** P < 0.01; *** P < 0.001; ****p<0.0001 (unpaired t tests). Data are representative of results from three independent experiments.

To further prove the above conclusion, we tested the effect of c-Jun knockdown and RBM39 on PRRSV proliferation. The results illustrate that it is obvious that RBM39 and siRNA-c-Jun alone can promote the proliferation of PRRSV (Fig. 7H, J). However, when the two were co-transfected, the promotion of PRRSV proliferation not only did not increase synergistically, but decreased, indicating that the promotion of PRRSV proliferation by RBM39 is through interacting with c-Jun (Fig. 7H, J).

In order to detect the effect of RBM39 on the phosphorylation level of proteins in the cell signaling pathway caused by PRRSV infection, 3D4/21 cells were transfected or co-transfected with PRRSV and RBM39, phosphatase Inhibitors/CIP was used to treat cell lysate supernatant and c-Jun antibody was used to detection for Western Blot analysis. In addition, phosphorylated antibody was used to detect the phosphorylation level of the protein 24 hours later. The results showed that c-Jun was heavily phosphorylated after PRRSV (Fig. 7D, K), which triggered subsequent immune responses. However, under the condition of RBM39 overexpression, the phosphorylation of c-Jun caused by PRRSV infection was greatly reduced and and in a dose-dependent manner (Fig. 7K, L). However, the c-Jun and phosphorylation level of JNK did not decrease significantly in the co-transfection group (Fig. 7G, K), indicating that RBM39 mainly does not reduce the phosphorylation of JNK triggered by PRRSV infection. In summary, the overexpression of RBM39 reduces the phosphorylation level of c-Jun after PRRSV infection, and thus down-regulates the AP-1 signaling pathway. This may be the mechanism by which RBM39 promotes PRRSV proliferation.

### PRRSV infection triggers nuclear export of RBM39 and c-Jun

To investigate whether PRRSV infection can cause the intracellular localization changes of RBM39 and c-Jun in 3D4/21 cells, the cells were respectively transfected with HA-RBM39 and Flag-c-Jun plasmids, 12 h after transfection, cells were infected with PRRSV at an MOI of 0.4. In the subsequent immunofluorescence analysis, we noticed that before PRRSV infects the cells, RBM39 or c-Jun proteins was localized in the nucleus (Fig. 6C, D). After the virus infection, we observed that both RBM39 and c-Jun proteins had moved from the nucleus to the outside of the nucleus (Fig. 6C, D) and co-localization (Fig. 6E). These results illustrate that PRRSV infection caused the translocation of RBM39 and c-Jun from the nucleus to the outside and this phenomenon may be related to the mechanism of RBM39 promoting PRRSV proliferation.

### RBM39 binds to PRRSV RNA

RBM24, which belongs to the same RNA-binding protein family as RBM39, has RNA-binding function and can bind to the 5’- and 3’-terminal ends of HBV RNA. Here the RBM39 binds to viral RNA were also identified. To verify whether RBM39 can bind to PRRSV RNA, 3D4/21 cells were transfected with HA-RBM39 or Flag-RBM39 plasmids and infected with PRRSV 12 h after transfection. After another 24 hours, the cells were collected and lysed with RIPA supplemented with Murine RNase inhibitor (40U/μl, Yeasen). Part of the supernatant was used for immunoblotting analysis (Fig. 8A, B), and the remaining supernatant was carefully added to HA/Flag-tags beads and combined overnight at 4℃. After RNA was extracted and reverse transcribed, it was amplified with corresponding primers, and a single band with the same size as predicted was recovered and sequenced (Fig. 8C, E and F). Through the comparison between SnapGene and DNAMAN, we found that we correctly amplified Nsp4, NSP5, Nsp7, Nsp10-12, M and N genes of PRRSV (Fig. 8D), indicating that RBM39 binds to PRRSV RNA.

**Figure 8.**
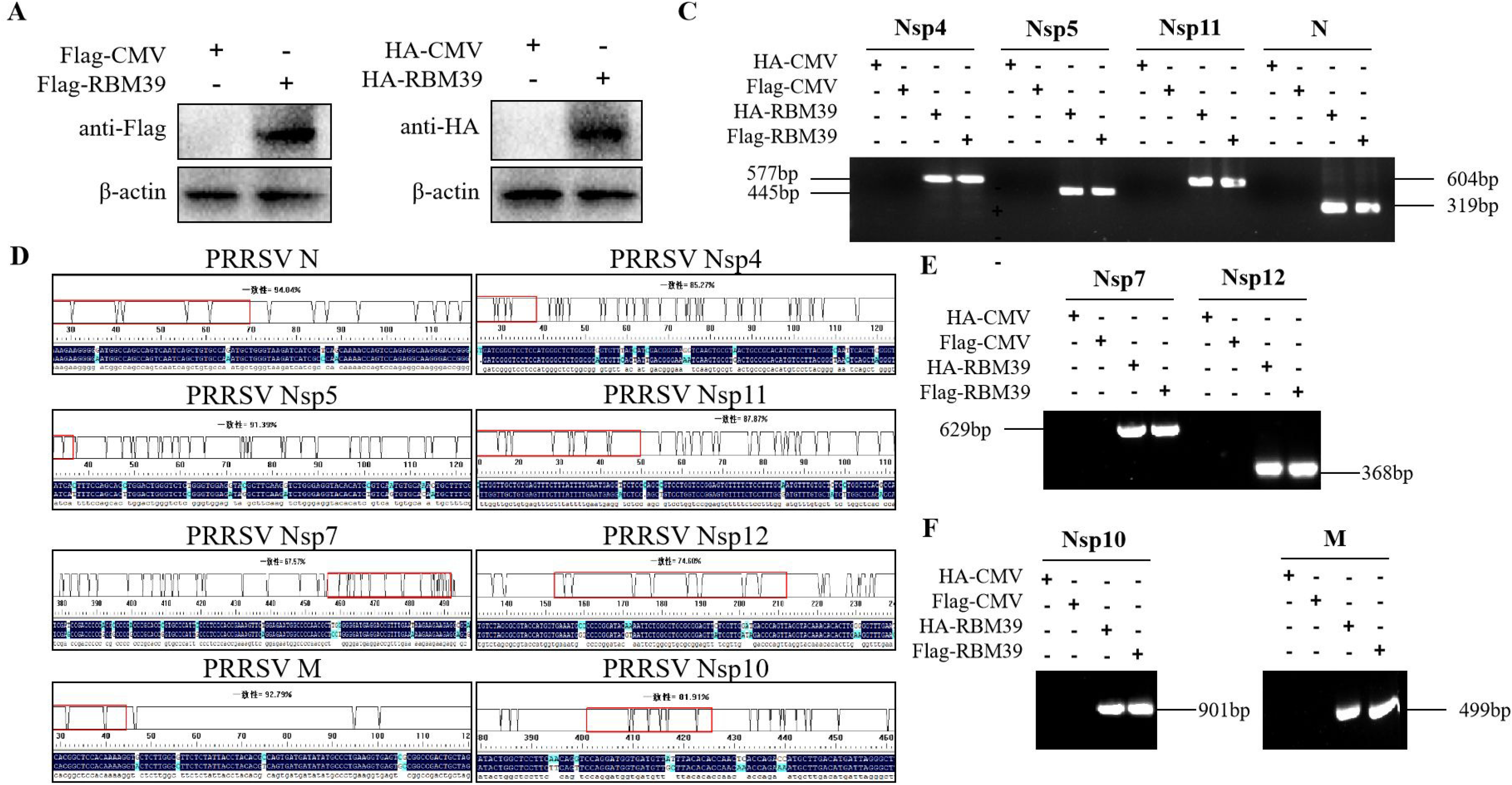
RBM39 binds to PRRSV RNA. (A and B) 3D4/21 cells were transfected with Flag/HA-RBM39 and infected with PRRSV 12 h after transfection. After another 24 hours, the cells were tested for immunoblotting analysis with anti HA/Flag mAb. (C, E, F) The agarose gel electrophoresis of PRRSV Nsp4, Nsp5, Nsp7, Nsp10-12, M and N PCR products after extracted RNA was reverse transcribed and amplified with corresponding primers. (D) Comparison of sequencing results using DNAMAN. Data are representative of results from three independent experiments.

## Discussion

RBM39 is an RNA-binding protein that has the function of binding RNA. In this study, we learned that RBM39 binds to PRRSV RNA and can promote its proliferation in 3D4/21 cells. According to reports, as the same RNA binding protein, RBM24 binds to the 3’-TR of HBV and stabilizes the stability of RNA(30). Therefore, we speculate that the upregulated cytoplasmic RBM39 promotes the combination of RBM39 with PRRSV RNA and further improve the viral RNA stability. In addition, the function of RBM39 in pre-mRNA splicing and its effects on the promotion of PRRSV proliferation should be further considered.

In summary, we report that porcine RBM39 negatively regulate the AP-1 signaling pathway and promote PRRSV proliferation. After PRRSV infection, both RBM39 and c-Jun showed phenomenon of translocation from cytoplasma to nucleus. The mechanism by which RBM39 promotes PRRSV proliferation can be summarized as follows. After PRRSV infection, the upregulated RBM39 reduces the level of phosphorylation of c-Jun by JNK and enhanced competitive combine of RBM39 with c-Jun and further prompt nucleocytoplasmic translocation of them to cytoplasm (Fig. 9). It is worth noting that RBM39 does not markedly affect the phosphorylation level of JNK and its upstream. Secondly, RBM39 competes with c-Fos to bind c-Jun and downregulate the formation of c-Jun/c-Fos complex, which further inhibits the transcription controlled by AP-1 promotor, and then down-regulates the activity of IFN- and other cytokines, and finally, promotes the proliferation of PRRSV. In addition, the enhanced binding capacity of RBM39 with PRRSV mRNA hints that the improved viral RNA stability also contribute to the PRRSV proliferation (Fig.9).

**Figure 9.**
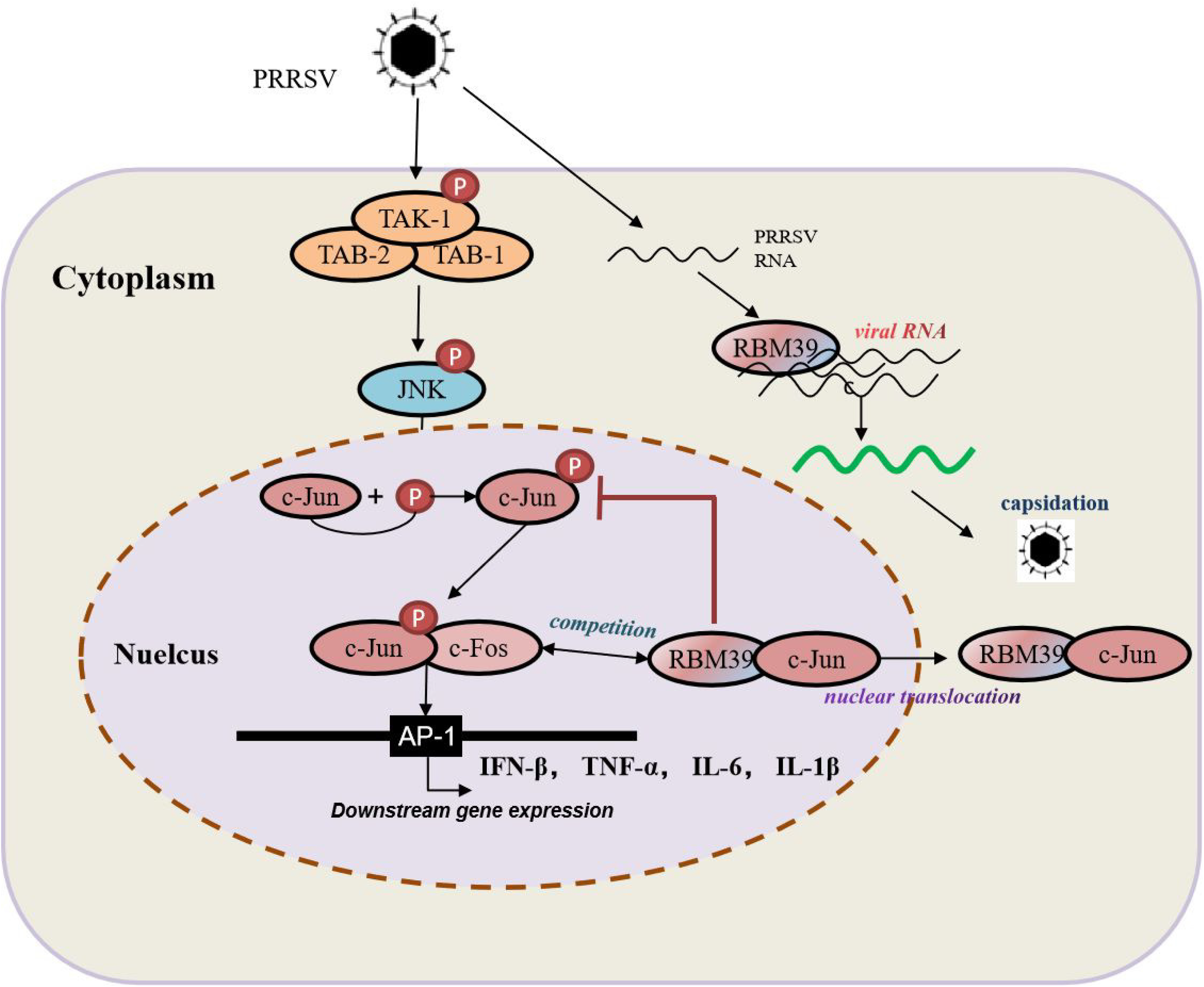
The diagram of the mechanism of RBM39 promoting PRRSV proliferation. After PRRSV infection, the combination of RBM39 and c-Jun translocates to the outside of the nucleus together under certain effects, which hinders the phosphorylation of c-Jun by JNK and reduces the level of c-Jun phosphorylation. At the same time, RBM39 competes with c-Fos to bind c-Jun and robs c-Jun of AP-1 dimer, which reduces the number of AP-1 dimers, and then down-regulates the activity of AP-1 pathway and its downstream signals, and finally, promotes the proliferation of PRRSV. In addition, the combination of RBM39 and PRRSV RNA may promote PRRSV proliferation to some extent.

The RNA-binding protein family can promote or inhibit the proliferation of viruses, and the mechanism of action is different. For example, Pseudouridine Synthase 4 (Pus4) could compete with the capsid protein (CP) of to bind to plus-strand Brome Mosaic Virus (BMV) RNAs and prevent virion formation(31). In higher eukaryotes, the production and expression of mature gene mRNA is a complex process that includes initiation of transcription, 5’-capping, splicing, and polyadenylation(32). RNA processing events play a vital role for specific transcription factors in pre-mRNA processing(33). Research have showed that RBM39 can regulated pre-mRNA processing in a steroid hormone receptor-dependent manner(32) and knockdown of RBM39 via siRNA changes the splice form of vascular endothelial growth factor A (VEGF-A)(32, 34, 35). Therefore, the mechanism of the precursor RNA splicing function of RBM39 and other RNA binding proteins after virus infection is worth exploring.

## Acknowledgments

This work was supported by the key underprop project of Tianjin Science and Technology Bureau in China (20YFZCSN00340) and the National Key Research and Development Program of China(2018YFD0500500).

## Author Contributions

Conceived and designed the experiments: JH H. Performed the experiments: YN S, XY L, YY G, RQ S, JX S, Z T, and LL Z. Analyzed the data: YN S and J H. Contributed reagents/materials: JH H. Wrote the paper: YN S and JH H.

